# Visualizing cellular heterogeneity by quantifying the dynamics of MAPK activity in live mammalian cells with synthetic fluorescent biosensors

**DOI:** 10.1101/760652

**Authors:** Min Ma, Pino Bordignon, Gian-Paolo Dotto, Serge Pelet

**Affiliations:** Department of Fundamental Microbiology, University of Lausanne; Department of Biochemistry, University of Lausanne

**Keywords:** MAPK signaling, single cell, fluorescent biosensor, live-cell imaging

## Abstract

Mitogen-Activated Protein Kinases (MAPKs) control a wide array of cellular functions by transducing extracellular information into defined biological responses. In order to understand how these pathways are regulated, dynamic single cell measurements are highly needed. Fluorescence microscopy is well suited to perform these measurements, however, more dynamic and sensitive biosensors that allow the quantification of signaling activity in living mammalian cells are required. We have engineered a synthetic fluorescent substrate for human MAPKs that relocates from the nucleus to the cytoplasm when phosphorylated by the kinase. We demonstrate that this reporter provides a better sensitivity relative to other similar biosensors and has allowed the monitoring of ERK MAPK activity pulses upon a single physiological EGF stimulation. In addition, we display its applicability to other MAPKs and in multiple cancer cell lines or primary cells as well as its application *in vivo* in developing tumors. Using our newly developed biosensors, dynamic single cell measurements with high temporal resolution can be obtained. These reporters can be widely applied to the analysis of molecular mechanisms of MAPK signaling in healthy and diseased state, in cell culture assays or *in vivo*.

## Background

Given that cells live in changing environments, they have developed complex biochemical networks to sense and transmit extracellular information. Intensive biological research has allowed the identification of multiple signal transduction pathways that relay the presence of nutrients, stress or hormones in the cell’s surroundings. The biochemical analysis of these pathways has allowed the identification of the major players implicated in these processes. However, to fully grasp the complex regulation of these biochemical signals, quantitative and dynamic measurements have to be performed.

An additional layer of complexity arises from the fact that each cell, even in an isogenic population, can display different cellular behaviors ^1^. Unfortunately, population-averaged measurements can prevent the identification of important cellular behaviors such as oscillations or bimodality ^2,3^. Therefore, the value of single cell measurements has been generally recognized. Microscopy and flow cytometry, as well as novel techniques such as mass cytometry or single cell sequencing, have gained in importance to unravel the biochemical signals taking place in cells. In the signaling field, the dynamics of the biological system is an essential parameter in the transmission of the biological information and the determination of cellular fate ^3,4^. Live-cell imaging is the only technique that can capture the temporal evolution of individual cells. However, more tools are required to enable the quantification of signaling activity in single cells in order to uncover novel biochemical regulations in signal transduction cascades.

In mammalian cells, quantification of kinase activity is of particular interest for signal transduction studies, because more than one third of cellular proteins have identified phosphorylation sites ^5^. Among these kinases, Mitogen-Activated Protein Kinases (MAPKs) play a central role in the signaling network of the cell. These proteins are evolutionary conserved serine/threonine kinases that mediate a wide range of cellular processes, including cellular growth, differentiation, adaptation to stress, apoptosis or motility ^6–9^. MAPKs can be activated by a wide range of extracellular signals. These environmental cues are typically detected by membrane sensors, which relay this information inside the cell to the MAPK signaling cascades. These cascades consist of three-core kinases (MAP3K, MAP2K, and MAPK) with sometimes an additional upstream MAP4K and a downstream MAPK-associate protein kinase. Once activated, the MAPK phosphorylates a wide array of substrates to modulate their activity. A minimal consensus site (SP or TP) is needed for MAPK phosphorylation, therefore the specificity of the process is achieved via specific protein-protein interaction domains. One conserved interaction motif is the docking site (DS), which is present in many substrates and activators of the MAPKs ^10–12^. In addition to the phosphorylation of substrates, MAPKs are tightly implicated in the induction of specific transcriptional program that can profoundly alter the cellular state.

In mammalian cells, three distinct groups of MAPKs have been described: Extracellular signal-Regulated Kinase (ERK), c-Jun N-terminal Kinase (JNK) and p38 ^13–15^. The analysis of targeted mutations in mice, and the development of specific inhibitors have begun to shed light on the function of these cascades in mammals. As MAPKs regulate a wide range of cellular processes, from proliferation to cell death, misregulation of MAPK signaling is implicated in the induction and progression of a broad range of diseases. Indeed, deregulation of the ERK pathway has been identified in approximately one third of all human cancers. The ERK pathway has been shown to be implicated in multiple fundamental cellular processes such as cell cycle entry, proliferation and differentiation ^16^. In this cascade, ligand-mediated activation of receptor tyrosine kinase triggers the Ras GTPase, which recruits and activates the MAP3K RAF. In turn, RAF phosphorylates and activates the MAP2K MEK, which finally activates the MAPK ERK. This kinase further relays the signal to many substrates. Among these substrates, there are several key transcription factors that will modulate the expression of genes implicated in cell fate decisions ^17,18^.

Due to the important role played by MAPKs in cellular fate determination and cancer progression, the ability to quantify their activity in live single cells is highly required. Numerous FRET bio-reporters have been developed and are commonly used to track the real-time kinase activities in single cells ^19–22^. To obtain a robust FRET signal, a large change in conformation between the active and inactive state of the molecule is required. Unfortunately, in practice these changes are often small and require a tedious optimization procedure to be improved ^23^. In addition, the FRET probes block two fluorescent channels of the microscope, therefore limiting their application to monitor multiple cellular activities within the same cell.

Due to the complexity in obtaining reliable FRET measurements, novel biosensors based on fluorescent protein relocation have been developed. One strategy is to modulate the shuttling of a fluorescent protein between the nucleus and the cytoplasm by phosphorylation. In mammalian cells, the KTR (Kinase Translocation Reporter) was developed to monitor ERK, JNK and p38 activities ^24^. In yeast cells, the SKARS (Synthetic Kinase Activity Relocation Sensor) has been engineered to monitor MAPK activity in the mating and cell wall integrity pathways ^25^. More recently, a new strategy based on protein aggregation has been presented ^26^, where the phosphorylation of a synthetic substrate by the kinase leads to the formation of bright fluorescent foci. One key advantage of these reporters is that they occupy a single fluorescent channel in the microscope. The combination of multiple relocation sensors within a single cell becomes feasible allowing the correlation of multiple signaling activities within the same cell ^24,27^.

In this paper, we have adapted the SKARS approach to quantify ERK, JNK and p38 activities in mammalian cells. We demonstrate that this sensor provides an improved readout compared to the KTR reporter by co-expressing both sensors in the same cells. Thanks to its sensitivity, this biosensor allows to monitor complex signal transduction dynamics in single cells that display a large heterogeneity. In addition, we show that it can be applied to a wide range of cell types *in vitro* and in developing tumors *in vivo*.

## Results

### The concept of SKARS biosensor for mammalian MAPK signaling

Nuclear Localization Signals (NLSs) are small amino acid sequences that confer to proteins the ability to be transported in the nucleus. These domains are highly conserved throughout eukaryotes, and for instance the SV40 NLS is commonly used in yeast as well as in mammalian cells. These NLS are positively charged peptides and often consist of a mix of four lysines or arginines. These positive charges enable a strong interaction with the importin, which will promote the transfer of the associated proteins through the nuclear pores into the nucleus ^28^. The amount of nuclear enrichment is directly linked to the ability of the NLS to bind to the importin ^29^. Point mutations in the NLS consensus sequence to uncharged amino-acids decrease the accumulation in the nucleus. Similarly, addition of negatively charged residues in the vicinity of the positively charged patch weakens the nuclear enrichment ^30^. These negative charges can be conveyed by phosphorylation events next to the NLS sequence. This strategy is often present in endogenous proteins were the nuclear enrichment of proteins is directly controlled by signaling events ^31–33^.

Our SKARS biosensors aim at mimicking this natural phenomenon. We express in the cell a synthetic substrate for a MAPK that can shuttle between the cytoplasm and the nucleus based on its NLS phosphorylation status. We combine two NLSs in order to promote a strong enrichment of the biosensor in the nucleus under basal conditions. Both NLSs bear two MAPK consensus phosphorylation sites (SP). The specific phosphorylation of the serines is controlled by the presence of a MAPK docking site (DS) placed nearby ^34,35^. These short peptides forming the docking modules are well known to steer the enzymatic activity of the MAPK towards specific proteins and exchanging these sequences between different proteins will alter the regulation by the MAPK ^10^. Finally, a fluorescent protein is functionalized with the DS and the NLSs to generate our reporter. In the absence of kinase activity, the fluorescence signal will accumulate in the nucleus thanks to the NLSs; while in the presence of an active MAPK, the phosphorylation of this synthetic substrate will result in its relocation into the cytoplasm (Figure 1a).

**Figure 1.**
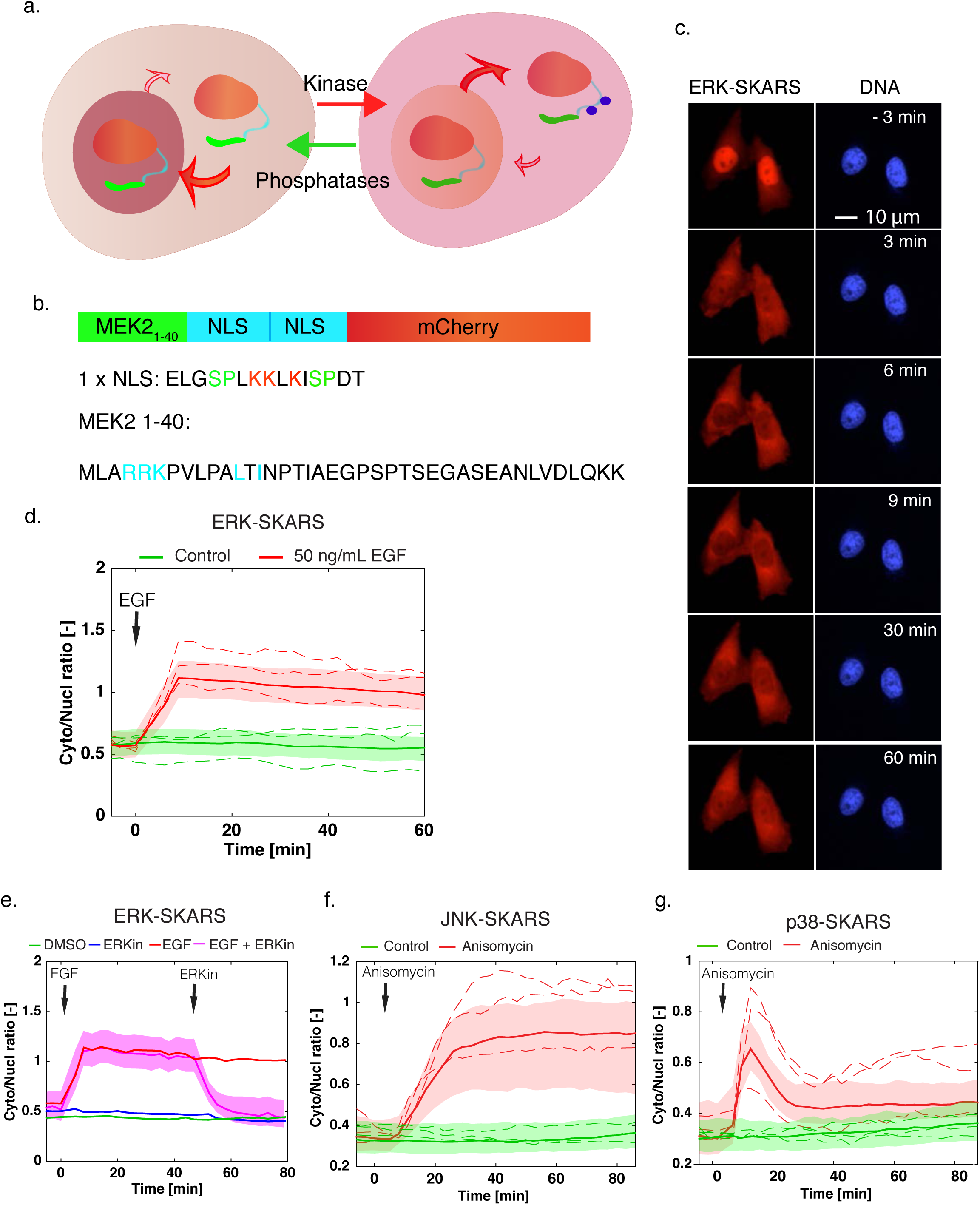
Principle and development of synthetic kinase activity relocation sensor (SKARS) to monitor MAPK activity in mammalian cells. a. Scheme of the SKARS relocation process. When the kinase is active, the NLS is functional and the sensor accumulates in the nucleus. When the kinase is active, it phosphorylates the SKARS, which diffuses into the cytoplasm. b. The ERK-SKARS contains three domains: the ERK docking site (MEK2 1-40), the two Nuclear Localization Sequences (NLS), and the fluorescent protein for visualization. Residues involved in interaction, nuclear import or phosphorylation are shown in cyan, red or and green respectively. c. Representative microscopy images of HeLa cells expressing the ERK-SKARS cells and stimulated with EGF (50 ng/ml) for 1 h period. In the red channel, the ERK-SKARS translocates from the nucleus to the cytoplasm. Nuclei are identified by a Hoechst staining. d. After quantification of the time-lapse movies, the ratio of the average cytoplasmic fluorescence of the average nuclear fluorescence (Cyto/Nucl ratio) is plotted as function of time. For all similar figures in this paper, the solid lines represent the median of the cell population and the shaded area the 25 and 75 percentiles of the population. The dotted lines represent a few single-cell traces extracted from the cell population. More than hundred single cells measured were plotted in the graph. The red curve represents HeLa cell treated with 50 ng/ml EGF (Number of cells: Nc = 340) and the green curve, mock-treated control cells (Nc = 449). e. Cyto/Nucl ratios of HeLa cells expressing the ERK-SKARS exposed to EGF stimulation (50 ng/ml) and ERK inhibition (FR 180204, 50 ng/ml) are plotted as function of time. EGF and ERK inhibitor were added at the time points indicated by the arrows. f. and g. HeLa cells expressing the JNK-SKARS (f) and the p38-SKARS (g) were treated with (red) and without (green) Anisomycin (50 ng/ml). The Cyto/Nucl ratio is plotted as function of time.

### Quantification of endogenous ERK activity by the ERK-SKARS

In order to adapt our previously developed yeast SKARS biosensor to report on mammalian MAPK activity, only the MAPK specific interaction module had to be changed. The 2xNLS peptide and the fluorescent proteins remained identical. Remarkably, the ability to use the same functional part of the sensor strongly suggests that this reporter could be applied to a wide range of eukaryotes. The docking site, however, needs to be adapted for each specific MAPK to be monitored due to ensure the specificity of the interaction. In mammalian cells, these interaction domains have been extensively studied and candidate DS motifs are relatively straightforward to identify from the literature. In order to target the ERK pathway, we selected a DS for ERK2 on its direct upstream activator MEK2, using the first 40 amino acids of this protein (Figure 1b) ^35^.

The ERK-SKARS was cloned into a lentivector and lentiviruses were produced to infect HeLa cells to generate cells stably expressing our construct. As shown in Figure 1c, the inactive form of the sensor accumulates in the nucleus of HeLa cells incubated in starved medium. Upon EGF stimulation (50 ng/ml), the ERK signaling cascade is activated and the MAPK can phosphorylate the serines in the vicinity of the NLS on the sensor. This leads to a rapid redistribution of ERK-SKARS to the cytoplasm. As illustrated in Figure 1c, six minutes after the stimulus, the nuclei of the two cells from the picture are strongly depleted from the bioreporter, providing a qualitative readout of MAPK activity in these cells.

To obtain statistically significant datasets, time-lapse movies were automatically quantified and single cell tracked over time using our image analysis platform ^36^. A nuclear marker (Hoechst) was used to automatically segment the nuclei, allowing to quantify the fluorescence intensity of the SKARS in the nucleus. The intensity in the cytoplasm is extracted from a ring surrounding the nucleus. The ratio of the cytoplasmic over nuclear intensities in each single cell is used as a proxy for MAPK activity. In Figure 1d, the median and 25-, 75-percentiles of the population response are plotted for more than 300 single cells stimulated with 50 ng/ml EGF and for an unstimulated control (PBS). Before stimulation, the ERK-SKARS is enriched in the cell nucleus, therefore the cytoplasm to nucleus ratio (Cyto/Nucl) is low (below 1). After EGF addition, the ratio increases suggesting that the sensor has been phosphorylated by active ERK in response to the EGF stimulation, leading to its shuttling from nucleus to the cytoplasm. The ratio remains high for the 60-minute duration of the time lapse (Figure 1 c and d). This sustained activation of ERK signaling upon high EGF treatment is also observed by Western blot and immuno-fluorescence staining (Sup Fig 1). Together, our data suggest that our biosensor reports on ERK activity. Compared to Western blot and immuno- fluorescence imaging which are routinely used to quantify ERK activity, our approach provides a dynamic readout of kinase activity in live single cells.

### Specificity of the ERK-SKARS

To demonstrate the specificity of our SKARS readout, we performed a number of control experiments. First, the four serines present in the NLSs were mutated to either non-phosphorylable (NLS-4A) or phospho-mimicking mutants (NLS-4E) (Sup Fig 2a and b). As expected, the NLS-4A is constitutively enriched in the nucleus (low Cyto/Nucl ratio), while the NLS-4E is depleted from the nucleus (high Cyto/Nucl ratio). In addition, both mutants lost the ability to respond to the EGF stimulus, therefore confirming that the phosphorylation of the four serines is controlling the nuclear localization of the sensor. Second, the five key residues in the docking site were mutated to alanine (non-docking MEK2, MEK2_ND_) (Sup Fig 2c). In the same cells, we compared the response of a non-docking mCherry SKARS and a functional sensor with a GFP fluorophore. While the reporter bearing the DS of MEK2 displayed the expected relocation from the nucleus into the cytoplasm, the ND reporter localization remained unchanged upon the addition of EGF (Sup Fig 2d to f), indicating a clear specificity of the substrate for the kinase interaction and phosphorylation of the substrate. Interestingly, these experiments also demonstrate that, while mutations in the DS or the NLS alter the function of the sensor, the fluorescent protein can be exchanged without perturbing the ability of the reporter to relocate. This feature allows to rapidly adapt the SKARS to specific imaging requirements and enables the combination of multiple sensors in the same cells.

To further validate the specificity of the ERK-SKARS reporter, a direct selective inhibitor of ERK (FR180204^37^) was applied to block the phosphorylation activity of the MAPK. The addition of the ERK inhibitor 45 minutes after the activation of the pathway leads to a rapid translocation of the sensor back into the nucleus (Fig 1e). In comparison, ERK activity was sustained for more than 80 min in the control cells, which were stimulated with EGF but not treated with the inhibitor (Fig 1e). In addition, pretreatment of the cells with the same ERK inhibitor abolished the EGF stimulated translocation of the ERK-SKARS (Sup Fig 3a). In contrast, cells pretreated with JNK or p38 inhibitors displayed an unperturbed relocation of the ERK sensor upon EGF stimulation (Sup Fig 3b and c). Taken together, these results demonstrate that the ERK-SKARS can efficiently and reliably quantify ERK kinase activity in single live mammalian cells.

### JNK and p38 SKARS

All MAPKs target the same phosphorylation motifs. Specificity in these cascades is thus achieved via distinct protein-protein interactions. Therefore, the docking sites in ERK interacting partners are different than the ones for JNK or p38 ^35,38^. To construct sensors for JNK and p38 signaling cascades, docking sites present in downstream transcription factors of the two MAPKs (c-Mef2 for p38 and c-jun for JNK) were selected to replace the MEK2 docking sequence of the ERK-SKARS (Sup Fig 4). Using lentiviral vectors, we generated HeLa cell lines stably expressing a JNK-SKARS or a p38-SKARS. As expected, anisomycin stimulation resulted in a fast translocation of both sensors into the cytoplasm (Figure 1f and g, and Sup Fig 4). Interestingly, the dynamics of the responses were very different for the two kinases. While anisomycin induced a sustained activation of the JNK pathway, p38 signaling was only transiently activated for 30 minutes, in agreement with previous observations ^39^. This experiment demonstrates once more the robustness of the design and versatility of these SKARS sensors, allowing the modulation of their specificity simply by exchanging the docking site of the reporter. This opens the door for targeting other kinases with similar modes of action (for example cyclins) or adapting the reporter to other organisms.

### Comparison of KTR and SKARS for the quantification of kinase activity

The design of our sensors bears many similarities with the KTR system ^24^. Therefore, in order to compare the relative abilities of the SKARS and the KTR to quantify MAPK activity, we generated cells stably expressing both reporters (Fig 2a). Both sensors were made spectrally compatible with an ERK-SKARS based on an mCherry protein, while the ERK-KTR was coupled to an mClover protein (a GFP variant). These two constructs are based on two different ERK DS: MEK2 for the SKARS and ELK1 for the KTR. But the major difference between these reporters resides in the peptides that control the nuclear to cytoplasmic shuttling. Indeed, the KTR combines an NLS and an NES (Nuclear Export Signal). The kinase activity drives the export of the construct because the phosphorylation weakens the NLS and enhances the efficiency of the NES. In comparison, for the SKARS, only the nuclear import rate is modulated by the phosphorylation of the NLS, and the sensor relies on the natural diffusion through the nuclear pores to exit the nucleus.

**Figure 2.**
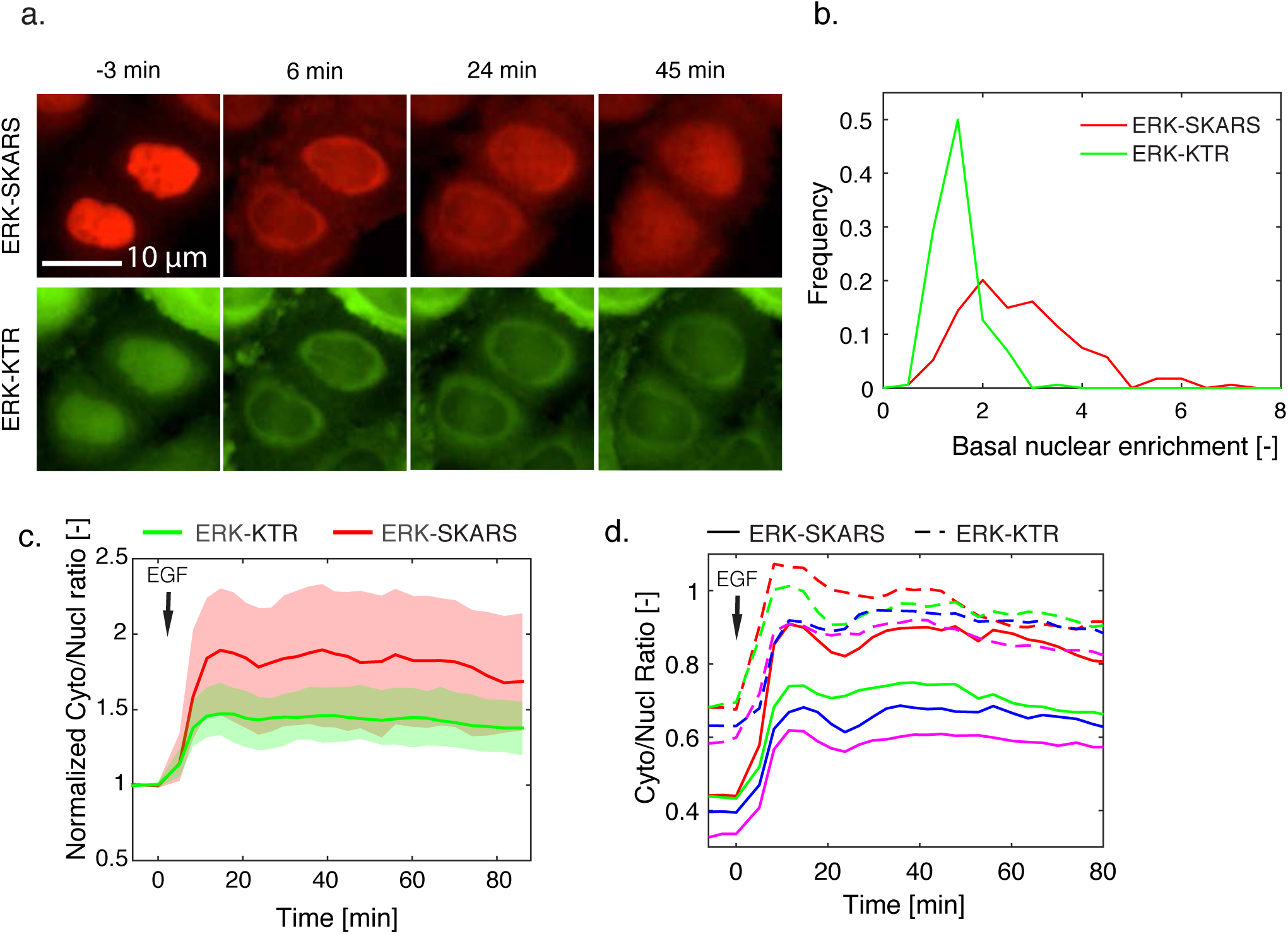
Direct comparison of ERK-SKARS and ERK-KTR in the same cells. a. Representative microscopy images of single HeLa cell carrying ERK-SKARS (red) and ERK-KTR (green) exposed to stimulation (EGF 50 ng/ml) and imaged at indicated time points. b. Histogram of SKARS and KTR nuclear enrichments before (Nucl/Cyto) of HeLa cells expressing ERK- SKARS (red) and ERK-KTR (green) before stimulation (mean of three first time points). c. Traces of Cyto/Nucl ratios from ERK-SKARS (red) and ERK-KTR (green) normalized to the basal level (the mean of the first 3 three time points) are plotted as function of time (Nc = 174). d. Single cell traces of the Cyto/Nucl ratios for ERK-SKARS (solid line) and ERK-KTR (dashed line) were plotted as function of time. Each color represents the measurements from one single cell, and both reporter traces for each cell is displayed.

To standardize both reporters’ expressions, we focused our analysis on cells expressing both reporters to intermediate levels to avoid sensitivity issues (Sup Fig. 5a). In these cells, the nuclear enrichment of the SKARS in basal condition is higher than for the KTR due to the reporter design (Fig 2b). In order to directly compare the responses of both sensors in the same cell, the single cell traces were normalized to their basal levels (mean of the first three time points before stimulus). As shown in Figure 2c, when cells were stimulated with EGF, both ERK-SKARS and ERK-KTR sensors could rapidly translocate from the nucleus to the cytoplasm, leading to a rapid increase of cytoplasm to nucleus ratio (Fig 2c). However, relative to the ERK-KTR population response amplitude (1.5 folds on average), the cytoplasm-to-nucleus ratio change is larger for the SKARS sensor (2 fold on average), highlighting a better signal-to-noise ratio for the later one. At the single-cell level, the traces for both sensors in the same cell were generally comparable but the amplitude of the SKARS response was generally larger (Fig 2d and Sup Fig 5b and c). Similar results were obtained when comparing the JNK-SKARS and the JNK-KTR in the same cells (Sup Fig. 6). Note that in the case of the JNK reporters, both sensors are based on the same docking site, thus the difference in relocation observed stems only from the phosphorylated part of the sensor. Because both techniques measure the ability of the MAPK to promote the translocation of the reporter out of the nucleus, starting with a higher nuclear pool of SKARS sensor provides a better dynamic range for the SKARS design compared to the KTR system.

### Single cell analysis of the dynamics of ERK activity

After validation of the ERK-SKARS reporter in mammalian cells, we next verified if we could observe heterogeneity in ERK activity between isogenic cells. In order to achieve this, we next isolated single-cell clones by serial dilution (See Methods). Three different cell lines bearing the ERK-SKARS were isolated and imaged. Since they displayed similar behaviors (data not shown), the data from only one of these lines is thus presented here.

First of all, we noticed that individual cells displayed a large variability of nuclear sensor enrichment before the EGF stimulation. This suggests that different basal activities of ERK are present in the individual cells of the population. Despite this large diversity, upon stimulation with a saturating concentration of EGF, the majority of the population displayed a strong ERK activation. In order to detect different activity patterns among the different cells’ responses, we first identified the non-responding cells (2% of the population), which displayed a weak relocation of the sensor (less than 0.05). The cytoplasm to nucleus ratios of the remaining cells were normalized between their basal value (average of the first three time points) and maximum values. K-mean clustering was applied to separate the time traces into four sub-populations. Figure 3a displays the response of almost 500 cells quantified in this experiment in a heat map. Each line corresponds to the normalized cytoplasm to nucleus ratio of a single cell. The traces were grouped in their individual clusters. Figure 3b displays the dynamics of ERK activation in the five sub-populations, with the normalized median response of each group.

**Figure 3.**
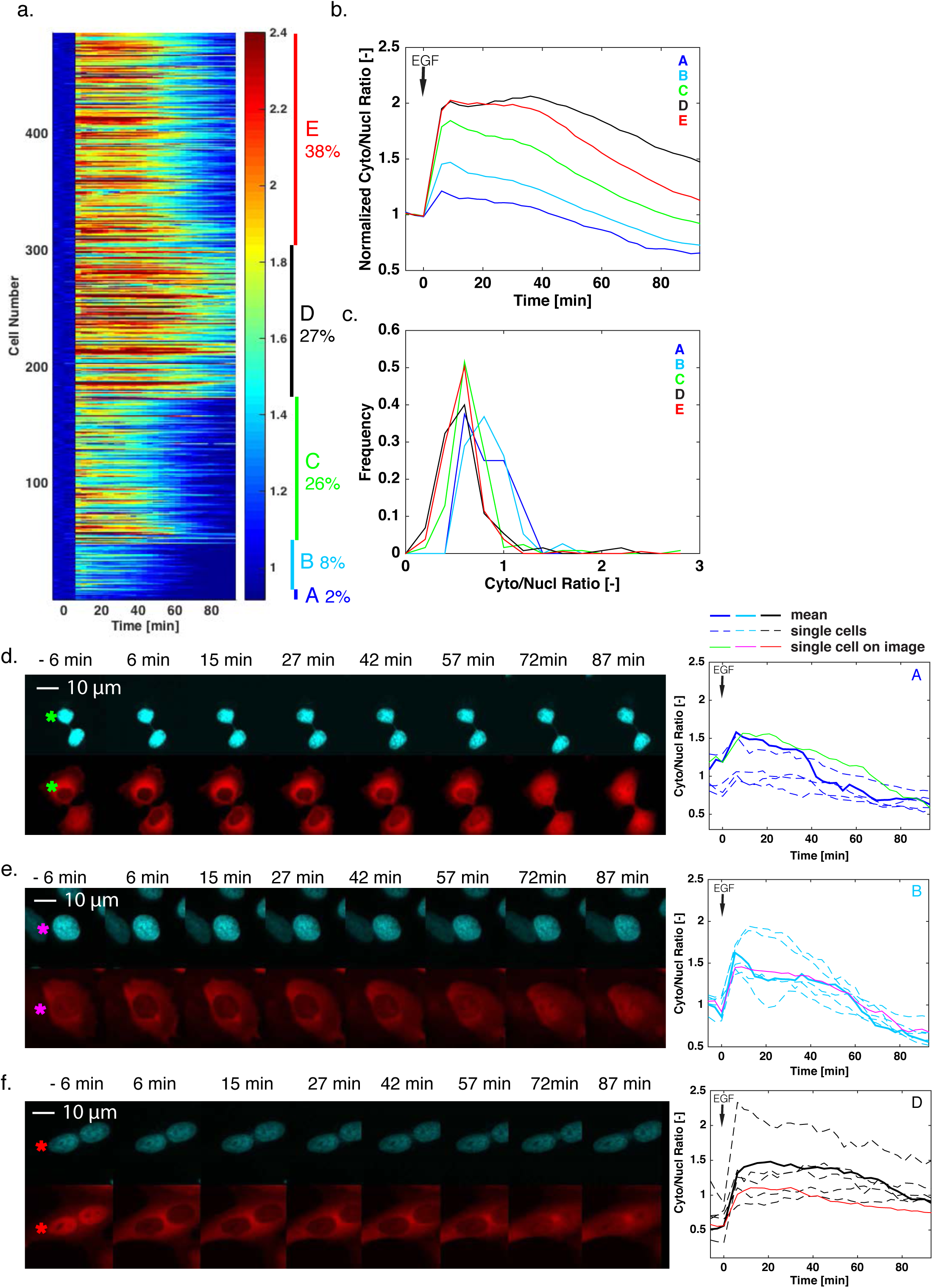
Cellular heterogeneity in ERK activity upon stimulation with high concentration of EGF. a. Heat map of the response of individual cells to a stimulation with EGF (50 ng/ml) in single clonal HeLa cells expressing the ERK-SKARS. Each line represents the cytoplasmic to nuclear ratio of one single cell normalized to the first three time points before stimulation. The cells were sorted using on k-mean clustering (See Methods). The different sub-populations are identified by the five colored bars on the right of the map. b. Median of the normalized Cyto/Nucl ratios from the indicated subgroups identified in the population. The legend indicates the relative prevalence of one cluster in the population. c. Histograms of the basal Cyto/Nucl ratios measured in the five subgroups. d., e. and f. Representative microscopy images of HeLa cells expressing the ERK-SKARS from three different sub-populations (A, B, D). The right panel displays the Cyto/Nucl measurements from six single cells from that sub-population. The solid line corresponds to the single cell identified in the microscopy images on the left with an asterisk. The dotted line represents the mean response from the sub-population.

Interestingly, the two sub-populations with the weakest responses (Fig 3, clusters A and B) display a higher basal cytoplasm to nucleus ratio (Figure 3c). This suggests that these cells already possess a substantial ERK basal activity and further activation of the pathway with EGF only leads to a minor increase in kinase activity. In the cluster A (non-responding cells, Figure 3d), we have observed an enrichment for cells that just came out of mitosis. This could be explained by a high ERK activity present during this cell cycle stage ^40^ or by an inability of the cells to enrich the sensor in the nucleus at this phase of the cell cycle. In the second cluster (Cluster B, Figure 3e), a weak activation of ERK is followed by deactivation. In these cells, the ERK activity level one and a half hour after the stimulus turns out to be lower than before the stimulus.

In the last three clusters (Clusters C, D and E, Figure 3f and Sup Fig. 7a and b), a strong activation of the pathway is observed. Interestingly, the differences in dynamics between these clusters arise mostly from a shoulder appearing at 40 min after the stimulus. Many cells in cluster D display a sustained activation that last longer than the 90 minutes of imaging, while in sub-population C and E the activity drops within the 90 minutes of the time-lapse movie. Since these cells were stimulated with a saturating level of EGF, we were expecting that the ERK pathway would display a high signaling activity and only minor differences between individual cells’ responses would exist. However, our data strongly suggest that the cell cycle stage and the basal activity of ERK can modulate the ability of the cells to transduce this extracellular information.

### Low doses of EGF induce a pulsatile response of the ERK pathway

We next wanted to assay the activity of the ERK pathway at more physiological concentrations of EGF ^41,42^. Our clonal cell line was imaged for more than 6 hours after stimulation with low doses of EGF. After a 0.5ng/ml and higher EGF stimulus, cells displayed a sustained activation of the pathway (Sup Figure 8a). At lower concentrations (from 0.1 ng/ml and lower), after displaying an initial peak, ERK activity dropped gradually. Interestingly, the median traces obtained from averaging more than 300 cells displayed small fluctuations. This feature arises from strong oscillations that take place in a sub-population of the cells. In order to better characterize this behavior, we imaged the cells with an improved temporal resolution and using EGF doses in the range where the oscillations are best observed (Figure 4a). The sub-population of cells that displayed oscillations was identified in the dataset by selecting peaks with an amplitude larger than 0.2 in the cytoplasmic to nuclear ratio of the sensor in single cell traces (Methods). At 0.1 ng/ml EGF, almost half of the population display at least three peaks (Figure 4b). At lower or higher concentrations, the oscillatory behavior tends to disappear and we cannot detect this type of oscillations in cells stimulated with 2.5 ng/ml or above. The heat map in Figure 4c represents the cytoplasmic to nuclear ratio of the sub- population of cells stimulated with 0.1 ng/ml where three or more peaks were identified. While the oscillations of individual cells can be visualized in this heat map, no global pattern emerges, indicating an asynchronous oscillation of the population. Indeed, when plotting individual single cell traces or looking at individual cells (Figure 4d and e, + Sup Movie 1), the strong pulses in ERK activity seem highly stochastic.

**Figure 4.**
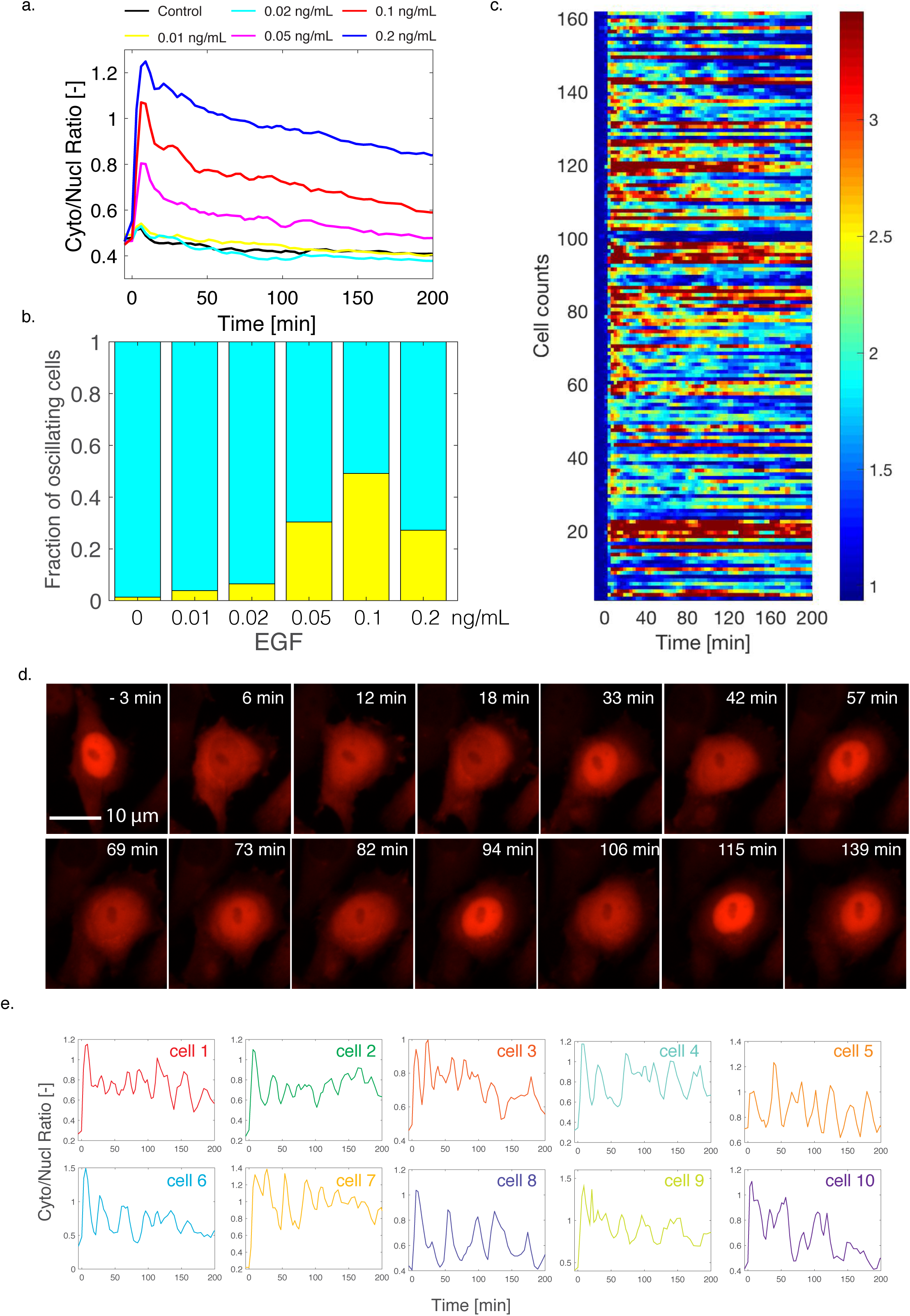
Monitoring of ERK activity pulses upon low doses of EGF. a. Median Cyto/Nucl ratio of ERK-SKARS in more than 400 single cells from a clonal HeLa cell population stimulated with low doses of EGF and monitored for more than 3 hours. b. Fraction of cells identified in the experiment from panel 4a that display three pulses or more in ERK activity. c. Heat map of the Cyto/Nucl ratio measured upon 0.1 ng/ml EGF treatment in HeLa cells. Only cells where three or more pulses were identified are displayed in this graph. d. Microscopy images of HeLa cell displaying pulses in ERK activation upon 0.1 ng/ml EGF stimulation. e. Various examples of single cell traces displaying fluctuations in ERK activation upon 0.1 ng/ml EGF treatment.

Multiple studies have uncovered oscillations in ERK activities ^43–46^. Some of these oscillatory patterns were observed with periods on the hour to multiple-hour time-scale and were linked to the proliferation of the cells. The behavior we observe is happening on the tens of minutes time scale. It displays many similarities to the ERK activation behaviors observed by Shankaran and co-workers using an ERK-YFP construct that relocates into the nucleus. However, they observed this pulsing behavior at 50 ng/ml EGF, a concentration at which we observe sustained activity of the pathway. Interestingly, this puzzling signaling pattern is a unique feature of the EGF response. Cells stimulated with FGF or PDGF did not display this oscillatory behavior (data not shown). From these preliminary data, we speculate that this pulsing behavior arises from an interplay between positive and negative feedback mechanisms impinging on the ERK pathway, however, more experiments would be required to understand the regulatory mechanism responsible for this behavior.

### SKARS in various cells types

All the above results show that the SKARS biosensors provide an efficient and reliable method to quantify the MAPK activity in real time in single HeLa cells. In the past few years, improved kinase reporters, such as KTR and FRET sensors, have provided the ability to quantify kinase activity in single cells. However, due to relative small dynamic change and high background noise, neither KTR nor FRET is sufficient to monitor kinase activity within a broad range of samples. Recently, the SPARK reporters demonstrated large dynamic changes allowing the monitoring of PKA and ERK activity in diverse samples ^26^. Although displaying a clear visual readout for MAPK activity, the quantification of kinase kinetics is more challenging due to a non-uniform spatial readout, which unfortunately hinders its application for the automated quantification of a large population of single cells. Here we show that, thanks to an improved readout, SKARS sensors can be used both to visualize and quantify the MAPK activity dynamics robustly in hundreds of single HeLa cells. In order to demonstrate the versatility of our sensor to other cell types, we tested the ERK-SKARS in various cell types, such as cancer cell lines (Figure 5a, 5b, 5c), primary cells (Figure 5d), neuronal cells as well as stem cells (data not shown). In the MDA-MB231 cancer cell line, we do not observe a response to EGF stimulus of the ERK-SKARS reporter because this cell line lacks HER2 (Human Epidermal growth factor Receptor 2) expression ^47^. Thus, EGF stimulation cannot induce ERK signaling activation (Fig 5c). In all other cell types tested, a strong and sustained response to EGF stimulus is observed and large heterogeneity in the single cell responses can also be monitored, as previously observed in the HeLa cells. These results confirm that the SKARS technique can be applied easily to different cell types, where ERK signaling dynamic ranges can vary significantly.

**Figure 5.**
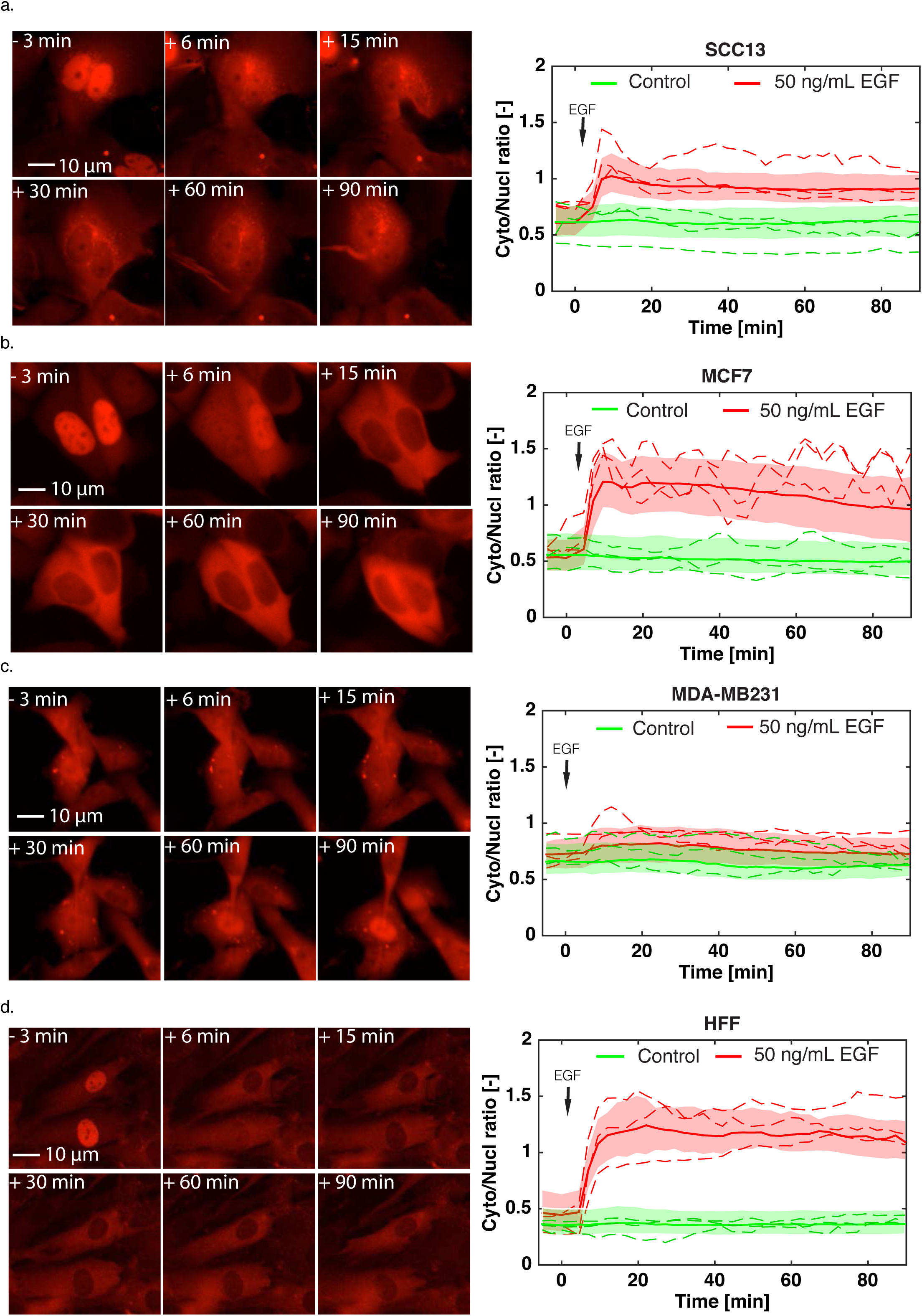
SKARS measurements performed in cancer and primary cell lines. a to d. Left panels show representative microscopy images of SCC13 (a), MCF7 (b), MDA-MB231 (c), HFF (Human Foreskin Fibroblast) (d) cells expressing the ERK-SKARS and stimulated with EGF (50 ng/ml). The right panels show the Cyto/Nucl ratio in the respective cell lines. Quantifications of the time-lapse movies for samples treated with EGF are plotted in red and control samples in green. For each trace, more than 100 single cells measurements were combined to generate the graphs.

### In vivo measurement of kinase activity with ERK-SKARS

Since misregulation of MAPK signaling has been observed in numerous diseases, especially in human cancers ^48,49^, a better understanding of cancer progression will undoubtedly require the ability to study signaling pathway activity in developing tumors. Currently, the most common way to quantify kinase activity in tissues remains immunohistochemistry. It is a complex and time-consuming method whose sensitivity is highly dependent on the specificity of the antibody and on sample preparation. Moreover, it only provides a snapshot measurement of signaling activity in the tissue. With the advent of novel microscopy techniques, it becomes feasible to image living cells directly in these complex environments ^50–52^. With their large dynamic range, SKARS reporters may enable the monitoring of kinase activity directly in tissues.

To test the application of the ERK-SKARS *in vivo*, we used a skin cancer orthotropic model, based on mouse ear intradermal injections of tumorigenic Squamous Cell Carcinoma (SCC) cells ^53^. We injected SCC13 cells expressing ERK-SKARS into the ears of 8- to 10-week-old female mice (Severe Combined Immuno Deficiency, SCID, CB17sc-m). After 14 days of growth, the cells formed a small tumor. The tissue was excised into 10-20 μm slides by Vibratome (Leica VT1200/S), stained with Hoechst and then imaged by confocal microscopy. Figure 6a shows the SCC13 cells (red) surrounded by the mouse-ear tissue (green). SCC13 cells displayed different levels of nuclear enrichment of the sensor *in vivo*, indicating that ERK is activated to different extents in these cells. In order to verify the activity of ERK in these settings, we performed another experiment where the SCC13 cells injected in the mouse ear were expressing a functional (MEK2_DS_) and a non-functional (MEK2_ND_) reporter (Figure 6b) as an internal control. In this situation, the lower nuclear enrichment of the ERK-SKARS in the GFP channel relative to the control in the RFP channel can provide a direct measure of kinase activity *in vivo*. Further experiments will be required to monitor the dynamics of MAPK activity in these settings; however, these results clearly show the versatility and robustness of SKARS reporters and open the window to a whole new field for cancer research.

**Figure 6.**
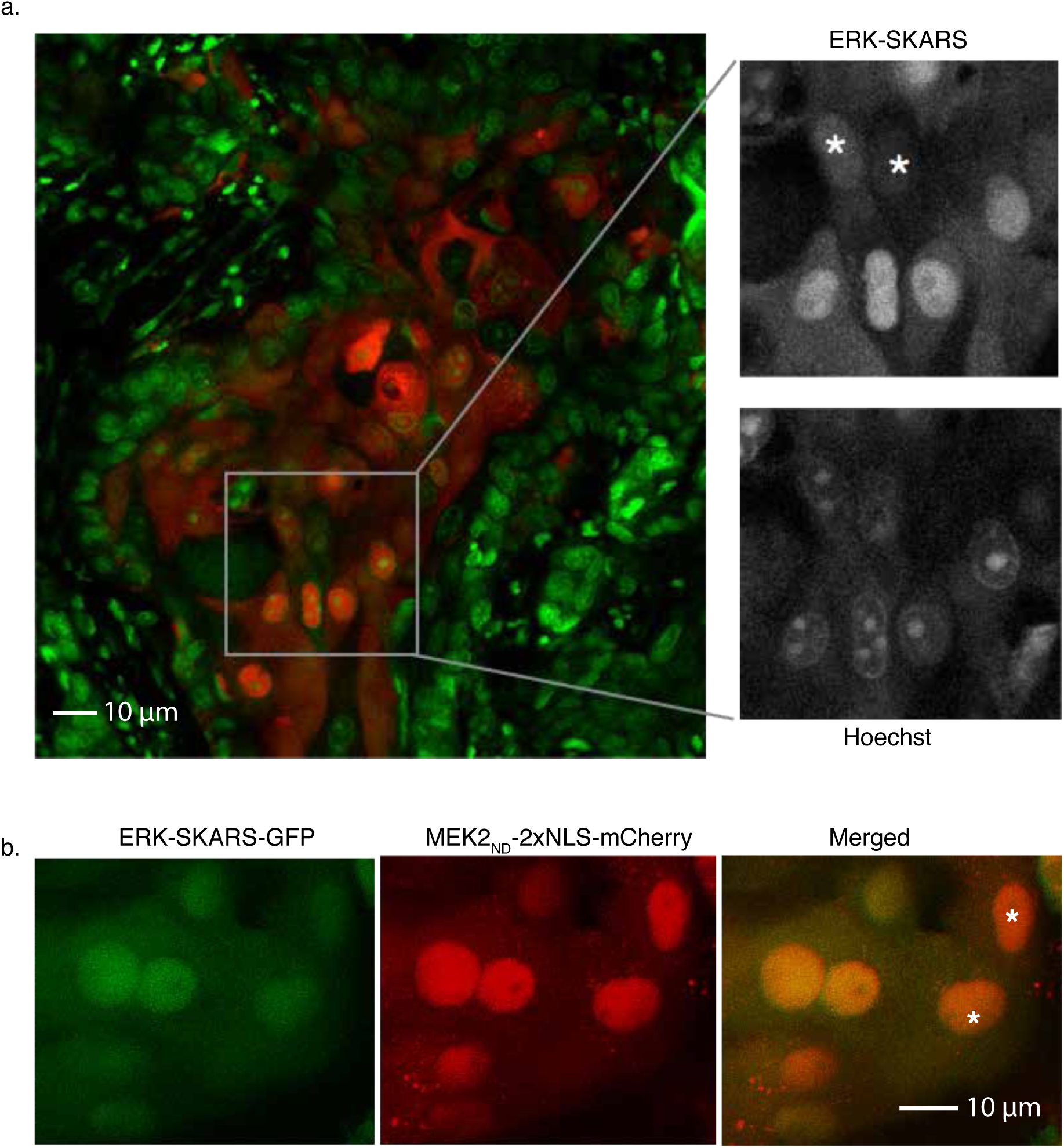
Application of SKARS for kinase activity quantification in tissue. a. Confocal images of HeLa cells expressing ERK-SKARS (red) in tumor implanted in a mouse ear. The green channel represents the nuclei of the cancerous and normal tissues, which were stained with Hoechst staining. The inset is a magnification of the image to display cells with different levels of nuclear accumulation of relative to the nuclear staining. Asterisk highlight cells with higher ERK activity. b. Confocal images of SCC13 cells expressing ERK-SKARS-GFP (MEK2DS, green) and MEK2_ND_-2XNLS-mCherry (red) in tumor implanted inside of mouse ears. Asterisk highlight cells with higher ERK activity.

## Discussion and Conclusion

The SKARS strategy provides a global measurement of kinase activity in a cell, while FRET biosensors can monitor local changes in kinase activity ^54,55^. For some studies, FRET biosensors might therefore have a clear advantage by adding a spatial information for signaling activity. However, in more difficult imaging conditions, like tissues, or when multiple sensors need to be quantified in parallel, SKARS can offer a better signal-to-noise ratio than other techniques. A second potential issue to be considered is that this reporter depends on the nuclear shuttling between the nucleus and cytoplasm. The efficiency of the nuclear import of the sensor can vary from cell to cell and can add undesired noise in the readout of the kinase activity. The addition of an internal control such as the non-docking construct could allow to correct for this artifact.

## Methods

### Animals

#### Mouse strain

NOD/SCID mice with interleukin-2 receptor gamma chain null mutation (Il2rg -/-) were maintained in the animal facility of the University Lausanne. All mouse work was carried out according to the Swiss guidelines and regulations for the care and use of laboratory animals, with approved protocol from the Canton de Vaud veterinary office.

### Human samples

Primary Human Foreskin Fibroblasts (HFF) were extracted from the discarded human tissue samples obtained from the Department of Pediatrics, at Lausanne University Hospital, with institutional approval and informed consent as part of institutional requirements.

### Cell culture

The cell lines and the primary cells were cultured and passaged in Dulbecco’s Mmodified Eagle’s Mmedium (DMEM) supplemented with 100 μg/ml streptomycin, 100 units/ml penicillin, 0.25 μg/ml amphotericin B (15240062, Gibco) and 10% heat inactivated Fetal Bovine Serum (FBS, 10270106, Gibco).

### Plasmids construction

All the SKARS plasmids for this study are constructed following to standard molecular biology protocol and the vector sequences were confirmed by exhaustively sequencing the cloned fragment. SKARS plasmids were constructed by cloning the docking site for the kinase of interest, the synthetic 2xNLS and the fluorescent protein of the sensor into a Lentiviral Vector backbone plasmid (pLV, a generous gift from Olivier Pertz, University of Bern).

### Cell line generation

Stable cell lines expressing the sensor of interest were generated by lentivirual transduction. Lentiviral vectors together with third generation packaging plasmids were transfected into 293T cells to generate lentiviruses. Packaged lentiviruses containing supernatants were added to recipient HeLa cells in the presence of 10 µg/ml Polybrene (AL-118, Sigma-Aldrich). At 3-5 days post transduction, cells were sorted based on fluorescent protein expression by flow cytometry (FACSAria III, BD Biosciences). After sorting, cells were cultured in DMEM. This medium is referred to as complete medium. The starvation medium is completed only with 0.5% FBS. Cells were cultured at 37°C in a humidified atmosphere containing 5% CO_2_. Singe-cell clones were isolated by serial dilutions of cells in a 96-well plate. To make sure that the colonies arose from single cells, the isolated single-cell colonies isolated have been recloned for a second time.

### Sample preparation and time-lapse imaging

The day before the time-lapse experiment, 10,000 cells/well of HeLa cells expressing the desired constructs were seeded onto the 96 well glass bottom optical imaging microplate (MGB096-1-2LG, Matrical Bioscience) coated with 10 µg/ml fibronectin (33010-018, ThermoFisher). 2 hours before the microscopy experiment, medium was replaced with Gibco FluoroBrite DMEM medium (A1896701, Gibco) supplemented with GlutaMAX, HEPES, sodium pyruvate and 0.5% FBS. For nucleus staining, we incubate the cells with 10 ng/ml Hoechst 33342 (H3579, Molecular Probes) for 1 hour.

Images were acquired on a fully automated inverted epi-fluorescence microscope (Ti-Eclipse, Nikon) controlled by micro-manager ^56^ using a 20X air objective and appropriate excitation and emission filters. The excitation is provided by a solid-state light source (SpectraX, Lumencor). The images were recorded with an sCMOS camera (Flash 4.0, Hamamatsu). A motorized XY-stage allowed recording multiple fields of view at every time point. GFP (50 ms), RFP (50 ms), DAPI (30 ms) and a bright-field (30 ms) images were recorded at time intervals varying from 2 to 3 minutes. During the experiments, the temperature (37°C), the CO_2_ (5%) and the humidity (95%) were kept constant using a temperature-controlled chamber and a CO_2_ controller (Cube and Brick, Life Imaging Service). Stimulations and chemical inhibitors were carefully added to the cells in the incubation chamber after the time-lapse imaging started. To activate ERK signaling, 50 ng/ml (or lower concentrations, where specified) EGF was added. To demonstrate the specificity of the SKARS-ERK, cells were pre-incubated with DMSO, MEK inhibitor (PD032591 100 nM), p38 inhibitor (p38 inhibitor 10 µM) and JNK inhibitor (JNK inhibitor VIII, 10 µM) for 30 min before imaging and stimulation. To inactivate the ERK signaling, ERK inhibitors were added 45 min after the EGF stimulation. To activate JNK and p38 signaling, Anisomycin 50 ng/ml was added to the cells.

### Data analysis

Time-lapse movies were analyzed with the YeastQuant platform ^36^. The cell nucleus and a 10-pixel-wide cytoplasm ring were segmented for each cell using the Hoechst image as a reference and quantified in each channel. The cell nucleus was tracked across all the frames of the movie. Multiple features of each cell were quantified. For each cell, the average nuclear intensity and the 10-pixel-wide cytoplasm ring in the fluorescent channel corresponding to the SKARS signal were extracted and used to calculate the cytoplasm to nucleus ratio providing a measure of kinase activity in each cell.

Data were further processed using Matlab software (R2017b, MathWorks). For quality control, among all cells tracked over the whole time-lapse experiment, only the cell traces with low variability in nuclear size and intensity were kept for further analysis. The basal level was calculated as the mean of the first three time points of the cytoplasm to nucleus ratio. To cluster the cellular responses in different sub-population, the non-responding cells were first identified by selecting cells with a change in Cyto/Nucl ratio smaller than 0.05. The other single cell traces were normalized between basal and maximal Cyto/Nucl ratio after stimulus. These normalized traces were fed to the *kmeans* function and the best clustering from 50 iterations was selected. In order to identify the pulses in the traces at low EGF concentrations, the single cell traces were passed to the *findpeaks* function. Only pulses (=peaks) with a minimal prominence of 0.2 were taken into consideration.

### Immuno-blotting (IB)

Total proteins were extracted with modified RIPA lysis buffer (50 mM Tris pH 7.4, 150 mM NaCl, 1 mM EDTA pH 8.0, 1% 1X NP40, 0.25% NA-deoxycholate, 2 mM Na-vanadate, 5 mM NaF, 1X protease inhibitors cocktail B (Santa Cruz, sc-45045)). Proteins were separated by SDS-PAGE gel, transferred onto PVDF membrane (Millipore), probed with primary antibody followed by HRP-linked secondary antibodies and detected by SuperSignal west pico chemiluminescent substrate (Thermo Scientific). The primary antibodies used for immnoblotting were rabbit anti-ERK1 Antibody (C-16) Santa Cruz sc-93 (1:1000), ERK2 Antibody (C-14) Santa Cruz sc-154 (1:1000), anti-GAPDH Antibody FL-335 Santa Cruz sc-25778 (1:1000), and anti-Phospho-p44/p42 MAPK (Erk1/2) Antibody (Thr202/tyr204) (D13.14.4E) XP Rabbit mAB (Cell Signaling, 4370). Anti-rabbit antibody (Rabbit IgG, HRP-linked whole Ab from donkey, Amersham NA 934) was used as a secondary antibody. After the detection of Phospho-ERK1/2, the membrane was stripped with stripping buffer (0.05 M Tris pH 6.8, 2% SDS, 0.8% beta-mercaptoethanol) for 30 minutes at 50 °C before detecting total ERK1 + ERK2.

### Immunofluorescence analyses (IFA)

10,000 cells were seeded onto a 13 mm glass coverslip in 24-well glass bottom plate coated with 10 ng/ml fibronectin. The next day, medium was replaced to starved medium and cultured for 1 hour, then the cells were stimulated for indicated times and fixed with 4% paraformaldehyde. After washing 3 times by PBS at room temperature, cells were permeabilized with 0.5% Triton X-100 in PBS for 10 minutes. Blocking of nonspecific epitopes was performed in blocking buffer (10% FBS in PBS) for 15 minutes at room temperature. The primary antibodies were applied at 1:100 dilutions in blocking buffer at 4 °C overnight. The fluorophore-conjugated secondary antibody (Donkey anti Rabbit IgG (H+L), Alexa Fluor 488 Thermo Fisher Scientific A-21206) and Hoechst 33342 were applied at 1:1000 dilutions in blocking buffer for 1-2 hours at room temperature in the dark. Images were acquired by Zeiss LSM 880 confocal microscope, and the images were processed with the FIJI software.

### Animal experiment: Intradermal ear injection and imaging

Mouse-ear injections of cells were carried out in 10-week-old male NOD/SCID mice with interleukin-2 receptor gamma chain null mutation (Il2rg -/-). SCC cells expressing the ERK-SKARS-mCherry or ERK- SKARS-GFP and MEK2_ND_-SKARS-mCherry were cultured in 10 cm dishes for 50-60 % confluence. After trypsinization and centrifugation, SCC13 cells were resuspended in 3 μl of sterile Hank’s Balanced Salt Solution, and injected intradermally in mouse ears through a 33-gauge microsyringe (Hamilton). Mice were sacrificed 3 weeks later for live tissue imaging analysis. Fresh tumors were sectioned into <1 mm slices and further analyzed through an inverted confocal microscope (Zeiss LSM880).

## Declarations

### Ethics approval and consent to participate

All mouse work was carried out according to Swiss guidelines for the use of laboratory animals, with protocols approved by the veterinary office of Canton de Vaud. All excised human samples were used with the approval of the University of Lausanne and CHUV. Informed consent was obtained as required.

### Consent for publication

Not applicable

### Availability of data and materials

The datasets generated and analyzed during this study will be made available upon request.

### Competing interests

The authors have no competing financial interests.

### Funding

Min Ma was supported by the Faculty of Biology and Medicine (FBM) interdisciplinary grant from the University of Lausanne. Work in the Pelet lab is supported by the Swiss National Science Foundation (SNSF) and the University of Lausanne. Work in the Dotto lab is supported by the SNSF. The funders had no role in study design, data collection and analysis, decision to publish, or preparation of the manuscript.

### Authors’ contributions

MM, SP and GPD conceived the study. MM performed the *in vitro* experiments. PB and MM performed the *in vivo experiments*. MM and SP analyzed the data. MM and SP wrote the paper.

## Acknowledgements

We thank all members of the Pelet, Martin and Dotto labs for helpful discussions, as well as Felix Naef and Nick Edward from the EPFL. Clémence Varidel and Marta Schmitt provided technical support. Olivier Pertz (University Bern) provided reagents. Mehdi Tafti and Cyril Mikhail (University of Lausanne) provided primary mouse neuronal cells.

## Supplementary Figure Legends

**Supplementary Figure 1.**
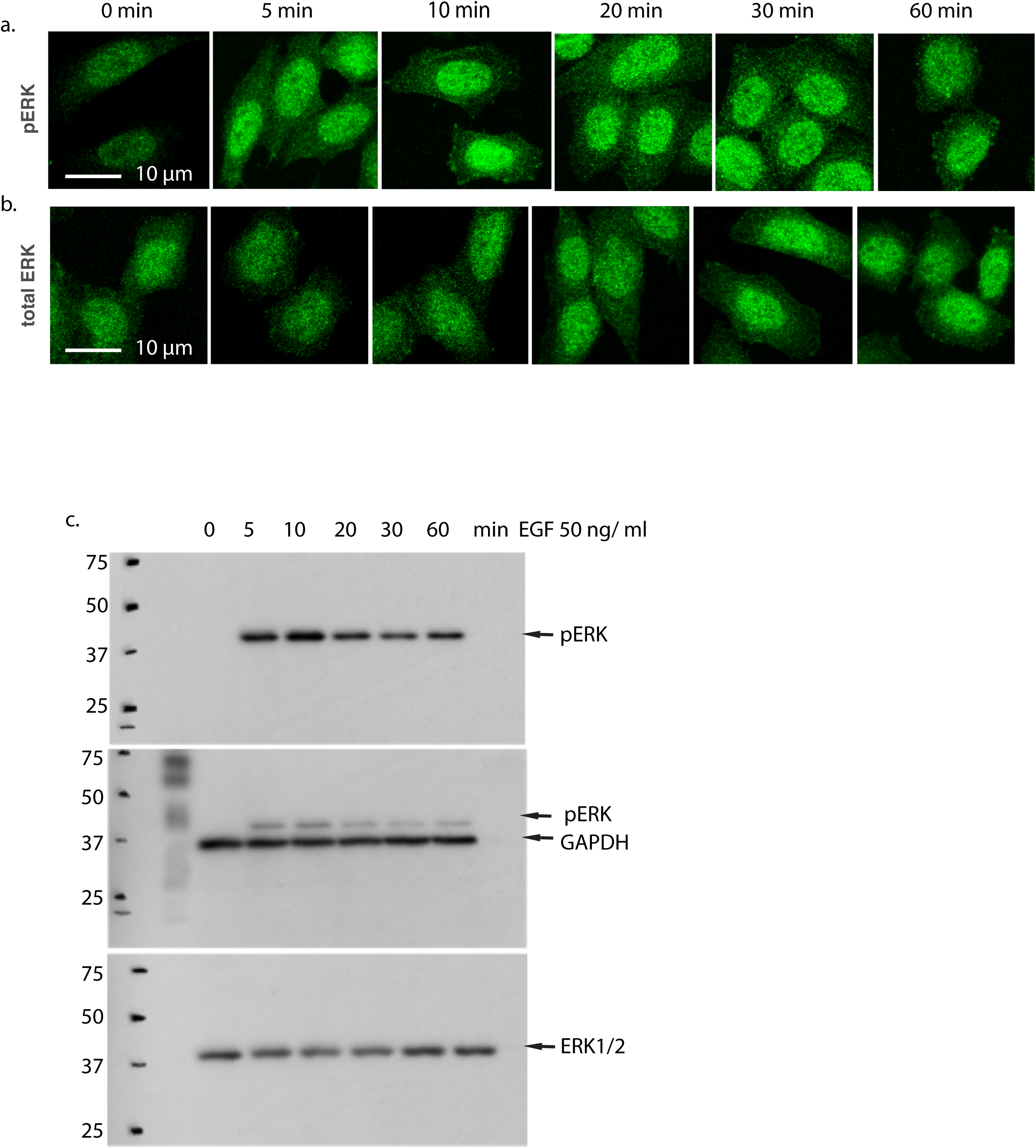
ERK activity measured by Western blot and immuno-fluorescence. a. and b. Representative immuno-fluorescence microscopy images of HeLa cells exposed to EGF (50 ng/ml) stimulation for the indicated period of time and probed for phosphorylated ERK (a) or total ERK levels (b). c. Time course analysis of total and phosphorylated ERK by Western Blot upon stimulation of Hela cells with 50 ng/ml EGF.

**Supplementary Figure 2.**
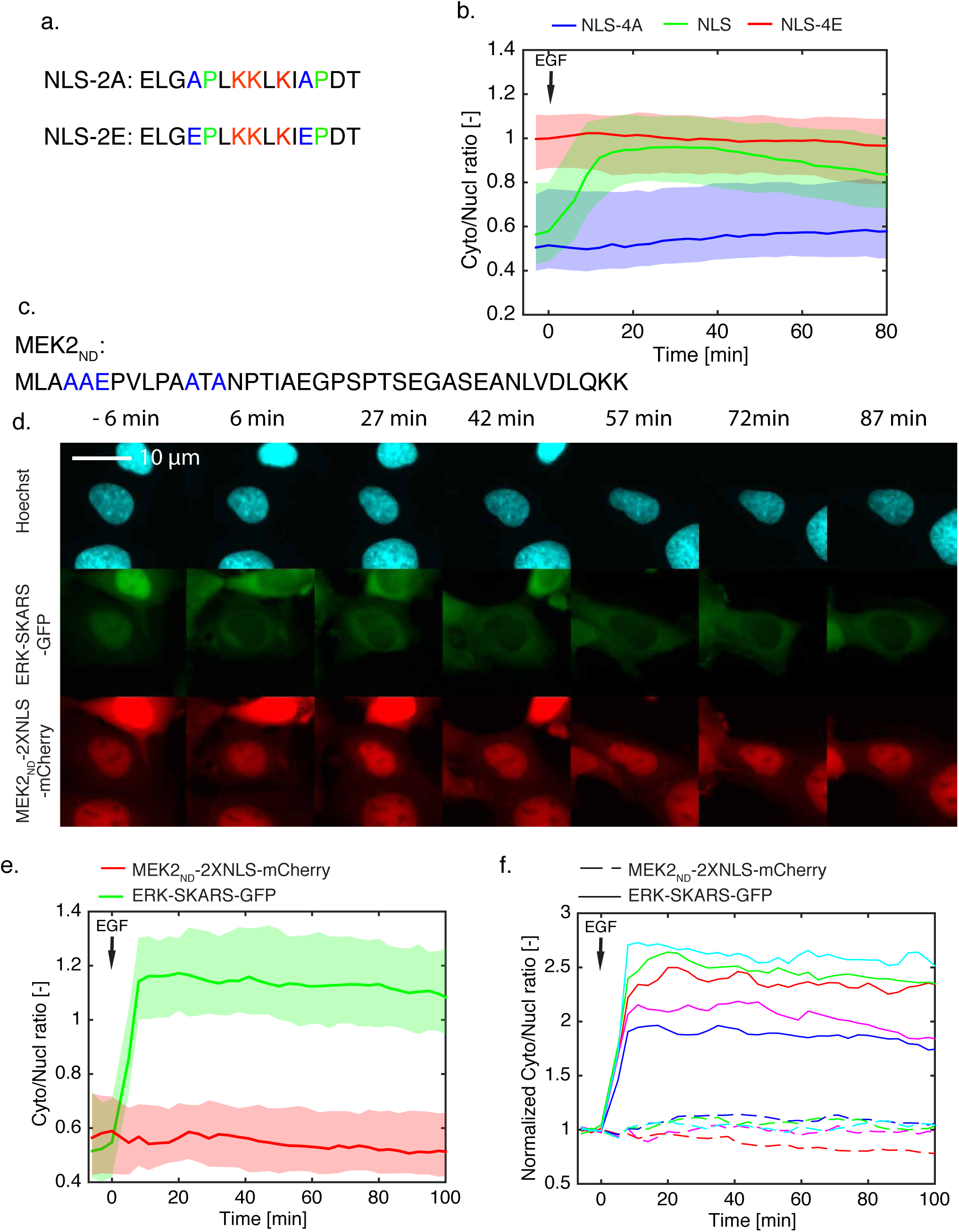
Characterization of the NLS and docking site specificity for ERK- SKARS function. a. Amino acid sequence of NLS-2A and NLS-2E mutation. b. Cellular translocation of ERK-SKARS requires phosphorylatable NLS. After quantification of the time-lapse movies, Cyto/Nucl ratio of HeLa cells is plotted as function of time. HeLa cells expressing ERK-SKARS (green), ERK-SKARS^4A^ (blue) and ERK-SKARS^4E^ (red) were stimulated with EGF 50 ng/ml. The solid lines represent the median of the cell population and the shaded area the 25 and 75 percentile of the population. More than 100 single cells were used to plot the graph. c. Amino acids sequence of the non-functional docking site of MEK2 for ERK. d. Microscopy images of HeLa cells co-expressing ERK-SKARS^DS^ (green) and ERK-SKARS^ND^ (red) exposed to EGF stimulation (50 ng/ml). HeLa cell nuclei were stained with Hoescht (cyan). e. Quantification of the time-lapse movie shown in d. The Cyto/Nucl ratio of the functional (green) and non-functional (red) sensors averaged over at least 120 single cells f. Few single cell measurements of the normalized Cyto/Nucl ratio. Each color corresponds to one single cell. For this cell, the solid line represents the ERK-SKARS^DS^ measured in the green channel and the dashed line represent the ERK- SKARS^ND^ in the red channel.

**Supplementary Figure 3.**
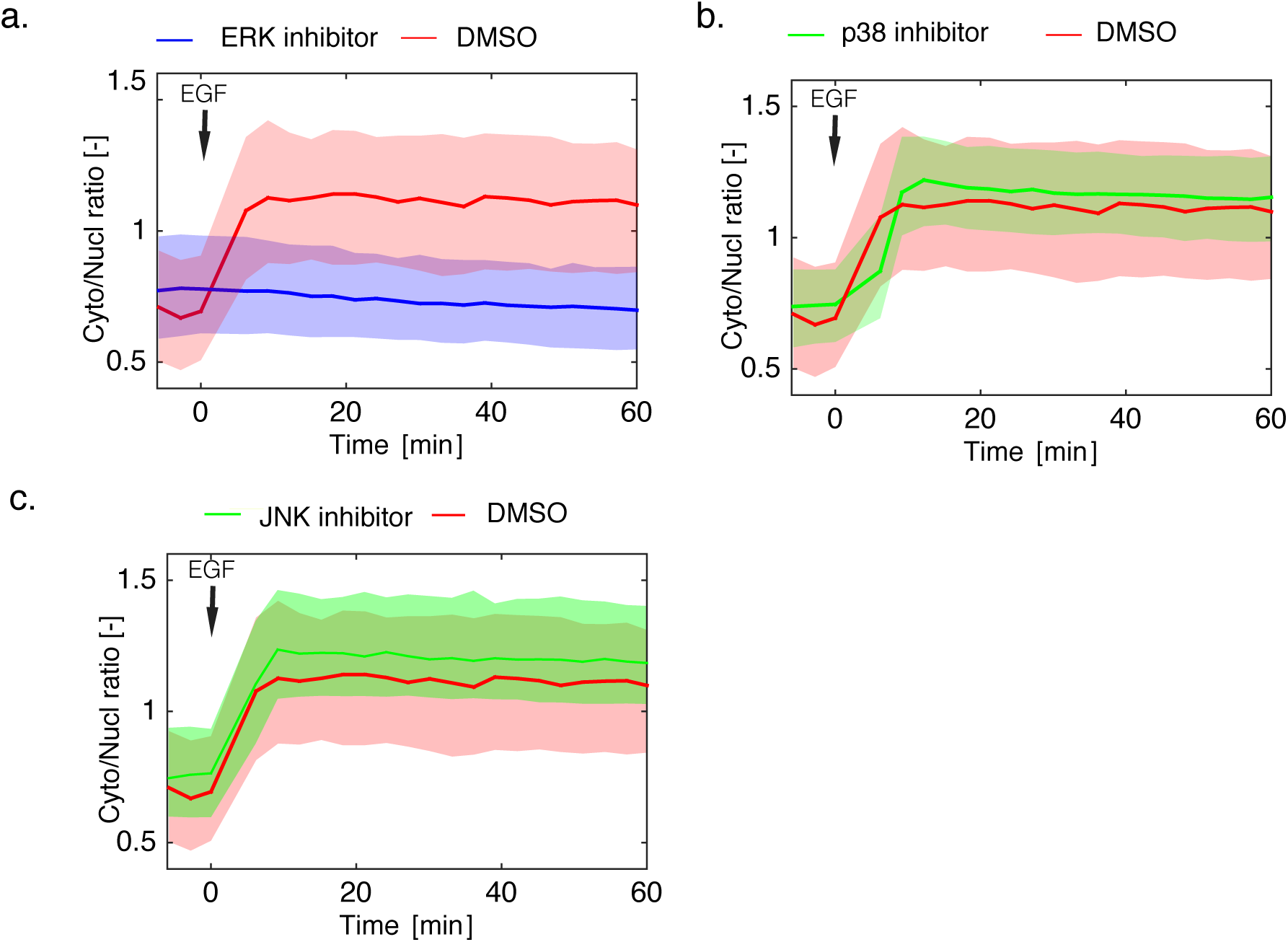
ERK-SKARS specificity validated using MAPK inhibitors. a. to c. HeLa cells expressing the ERK-SKARS were pre-incubated with DMSO (red), or ERK inhibitor (PD032591 100 nM, a, blue), p38 inhibitor (10 uM p38 inhibitor, b, green) and JNK inhibitor (10 uM JNK inhibitor VIII, c, yellow) for 30 min before imaging. Cells were subsequently stimulated with EGF 50 ng/ml.

**Supplementary Figure 4.**
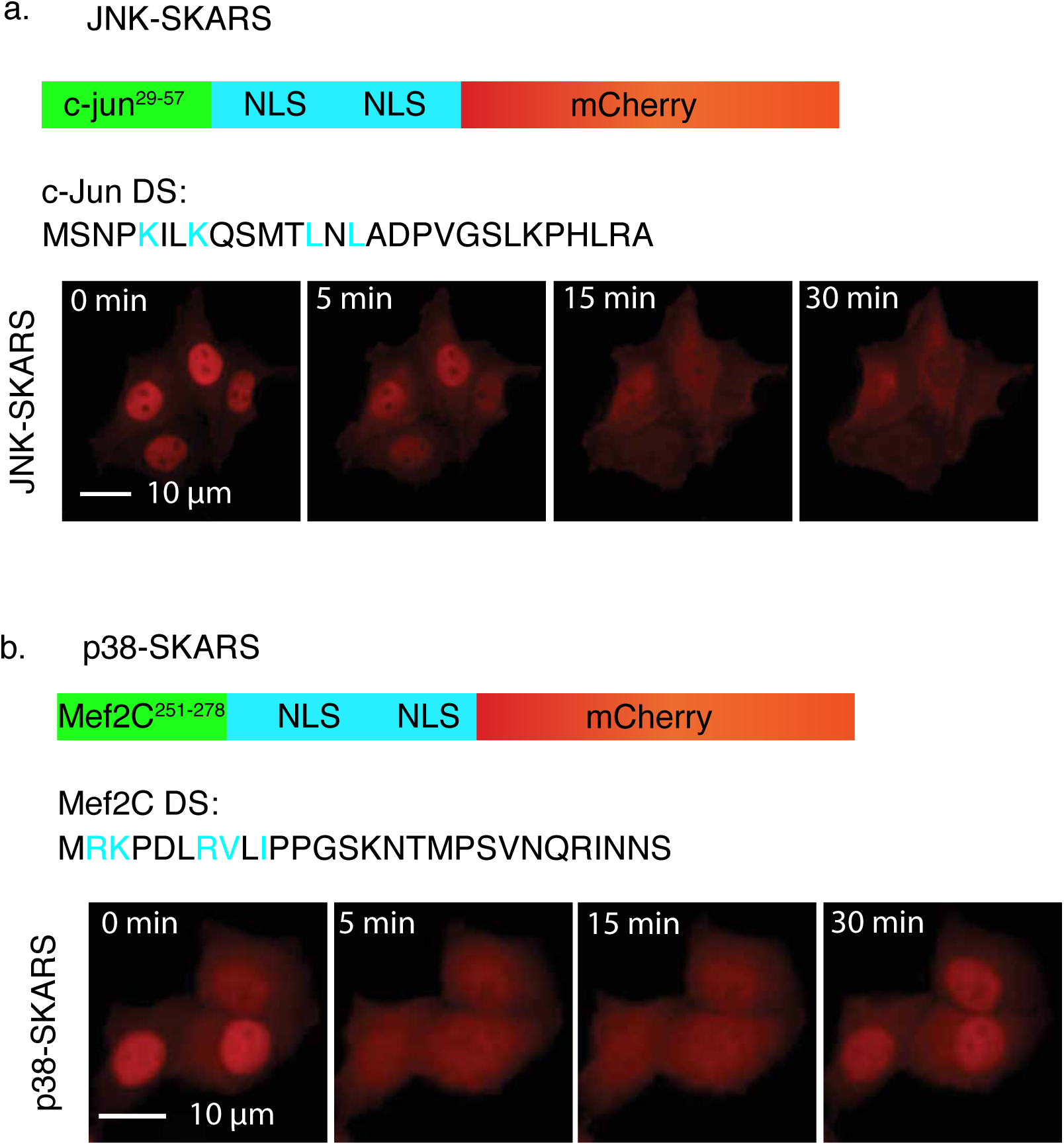
Images of JNK-SKARS and p38-SKARS sensors. a. Specific sequence used for developing JNK. The JNK-SKARS is composed of the c-Jun docking site (c-Jun, amino acids 29-57), double NLS and the mCherry protein. b. Microscopy images of HeLa cells expressing the JNK-SKARS and stimulated with Anisomycin (50 ng/ml). c. The p38-SKARS is composed of the Mef2C docking site (Mef2C, amino acids 251-278), double NLS and the mCherry protein. c. Microscopy images of HeLa cells expressing the p38-SKARS and stimulated with Anisomycin (50 ng/ml).

**Supplementary Figure 5.**
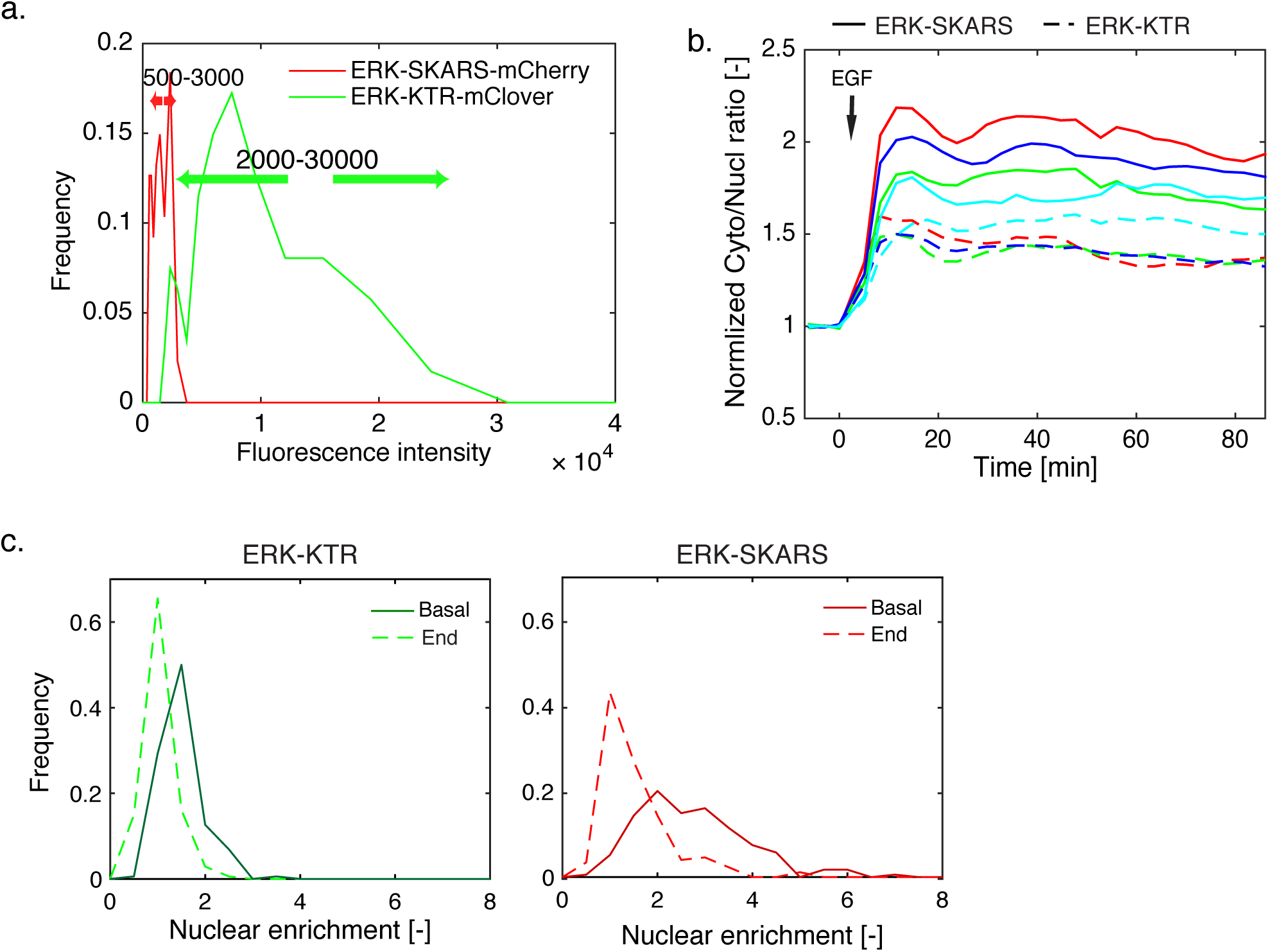
Comparison of the SKARS and KTR reporters. a. Comparison of the relative fluorescence intensities of the ERK-SKARS-mCherry (red) and the ERK-KTR-mClover (green). Cells expressing both sensors within the range of fluorescence intensities indicated by the arrows were kept for analysis. b. Few single cell measurements of the normalized Cyto/Nucl ratio following EGF (50 ng/ml) stimulation. Each color corresponds to one single cell. For this cell, the solid line represents the ERK-SKARS measured in the red channel and the dashed line represent the ERK-KTR in the green channel. c. Nuclear enrichment (Nucl/Cyto ratio) of the ERK- SKARS (red) and the ERK-KTR (green) before (solid line) and at the end of the time lapse (dashed line).

**Supplementary Figure 6.**
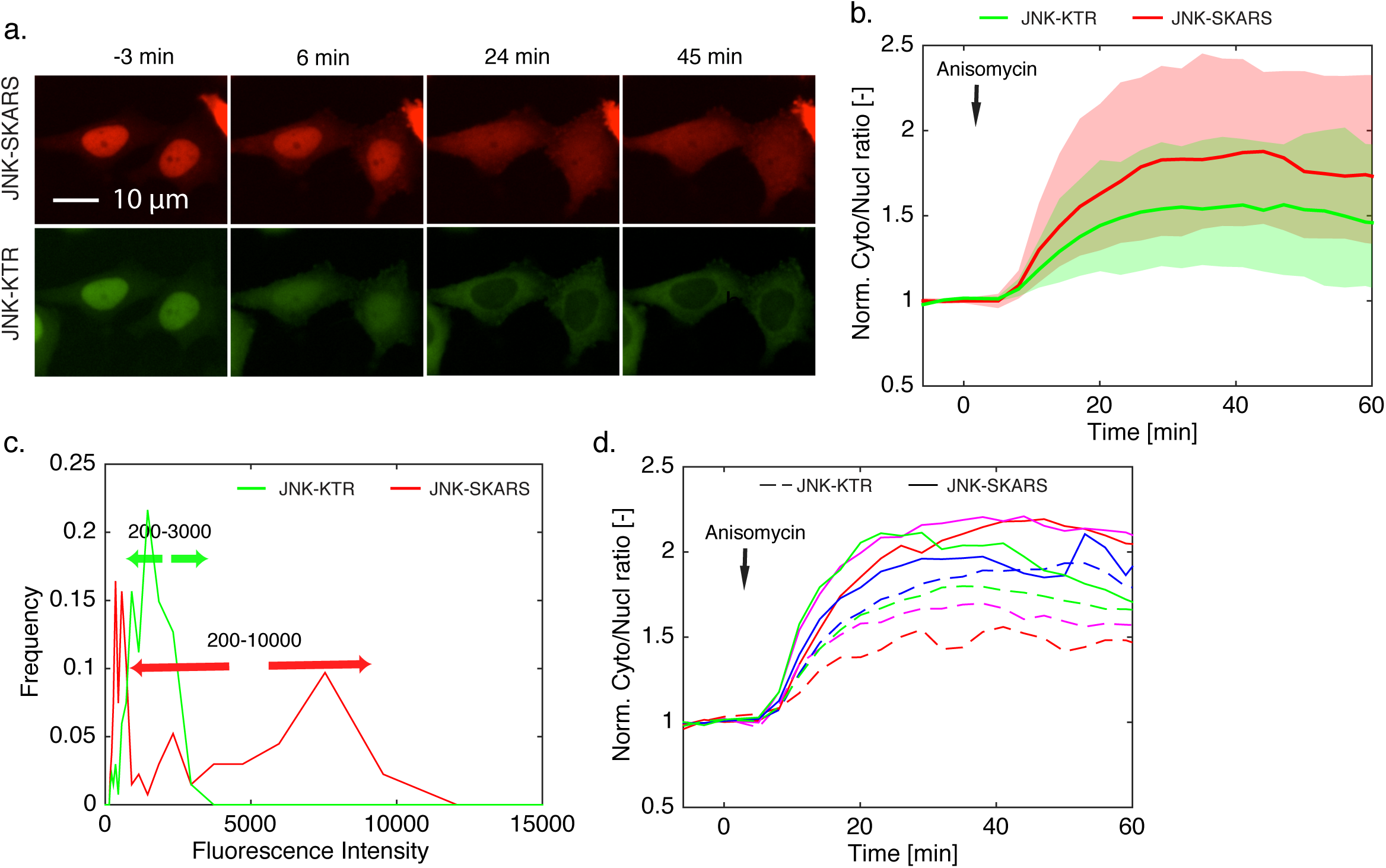
Comparison of JNK-SKARS and JNK-KTR. a. Microscopy images of HeLa cell carrying JNK-SKARS-mCherry (red) and JNK-KTR-mClover (green) exposed to Anisomycin (50 ng/ml) stimulation and imaged at indicated time points. b. After quantification of the time-lapse movies, traces of Cyto/Nucl ratio from JNK-SKARS (red) and JNK-KTR (green) normalized by the average of the first 3 time points are plotted as function of time. c. Histograms of the cell nuclear fluorescence intensity in the GFP (JNK-KTR) and RFP (JNK-SKARS) channels. 134 cells expressing both sensors within the range of fluorescence intensities indicated by the arrows were kept for analysis. d. Few single cell measurements of the normalized Cyto/Nucl ratio following Anisomycin (50 ng/ml) stimulation. Each color corresponds to one single cell. For this cell, the solid line represents the ERK-SKARS measured in the red channel and the dashed line represent the ERK-KTR in the green channel.

**Supplementary Figure 7.**
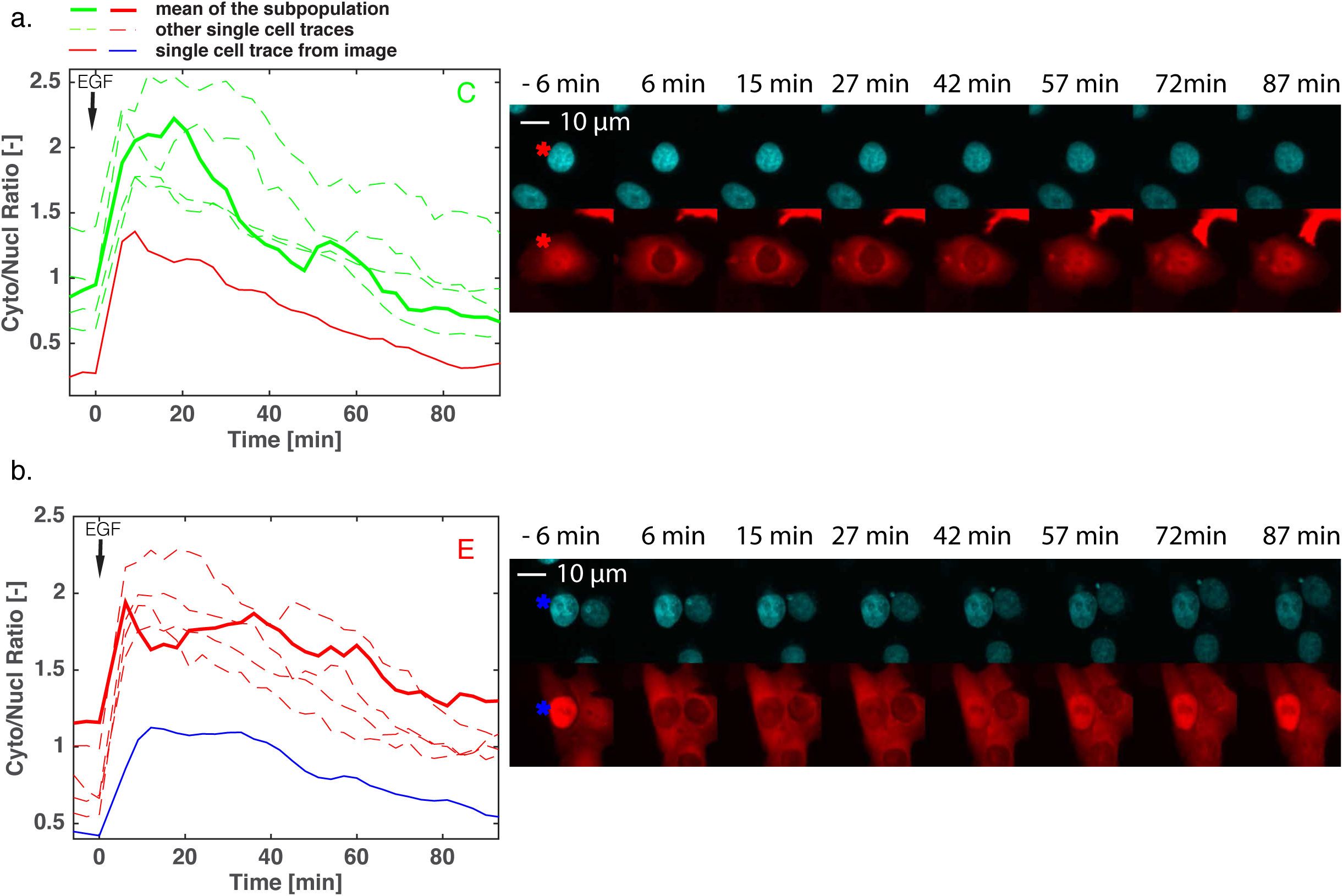
Single cell responses to high doses of EGF for two of the five clusters. a. and b. Microscopy images of HeLa cells expressing the ERK-SKARS from two different sub- populations (C and E). The right panel displays the Cyto/Nucl measurements from six single cells from that sub-population. The solid line corresponds to the single cell identified in the microscopy images on the left with an asterisk. The dotted line represents the mean response from the sub-population.

**Supplementary Figure 8.**
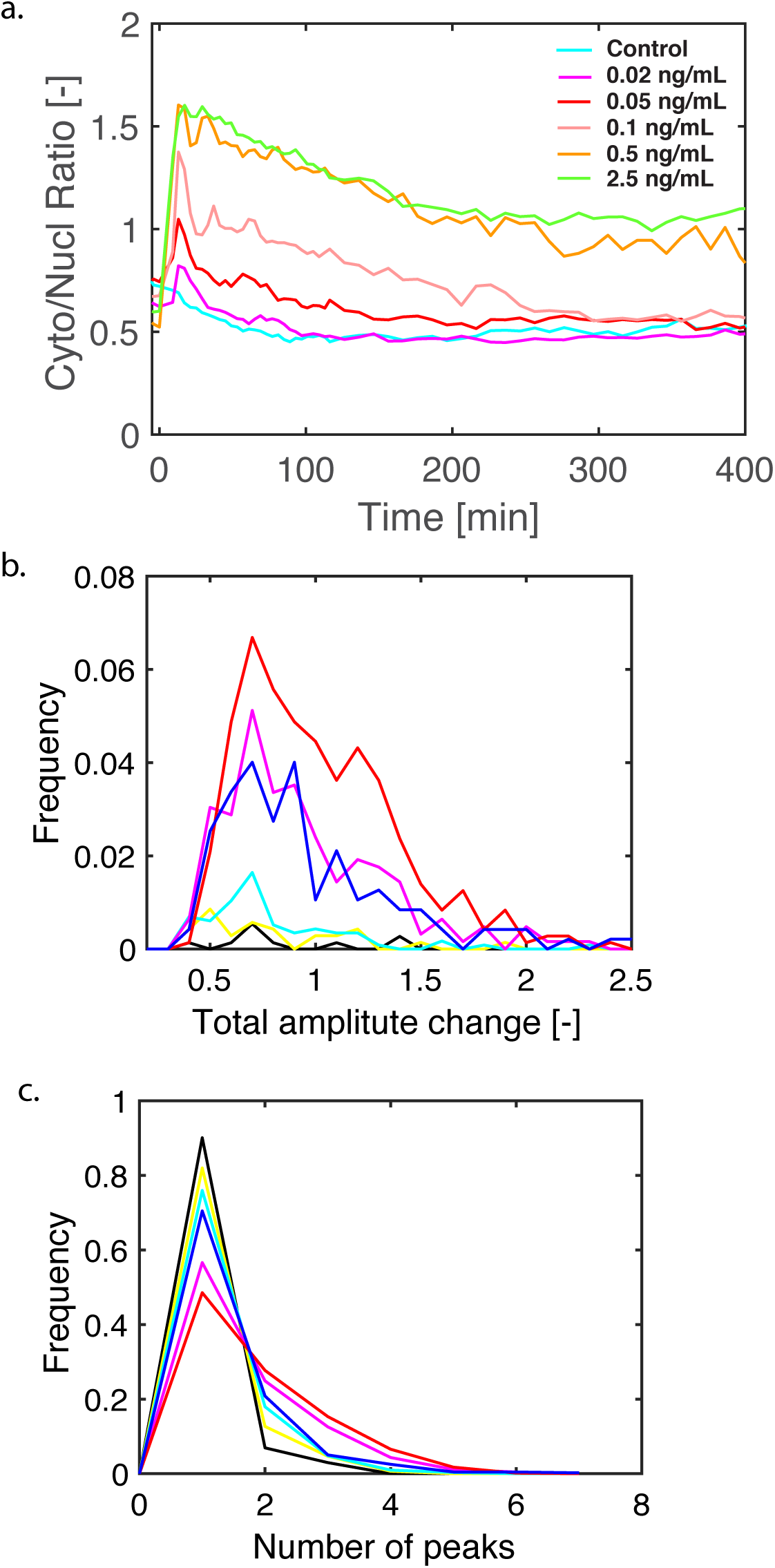
Analysis of pulses in single cell responding to low doses of EGF. a. Median Cyto/Nucl ratio of a clonal population of Hela cells expressing the ERK-SKARS responding to low concentrations of EGF (Nc > 300) for more than 6 hours. b. Histogram of the sum of the amplitudes of all the pulses in the traces displaying more than 2 pulses. The cell count in the histogram was normalized to the total number of cells measured in the dataset. c. Histogram of the number of peaks detected at the different concentrations of EGF.

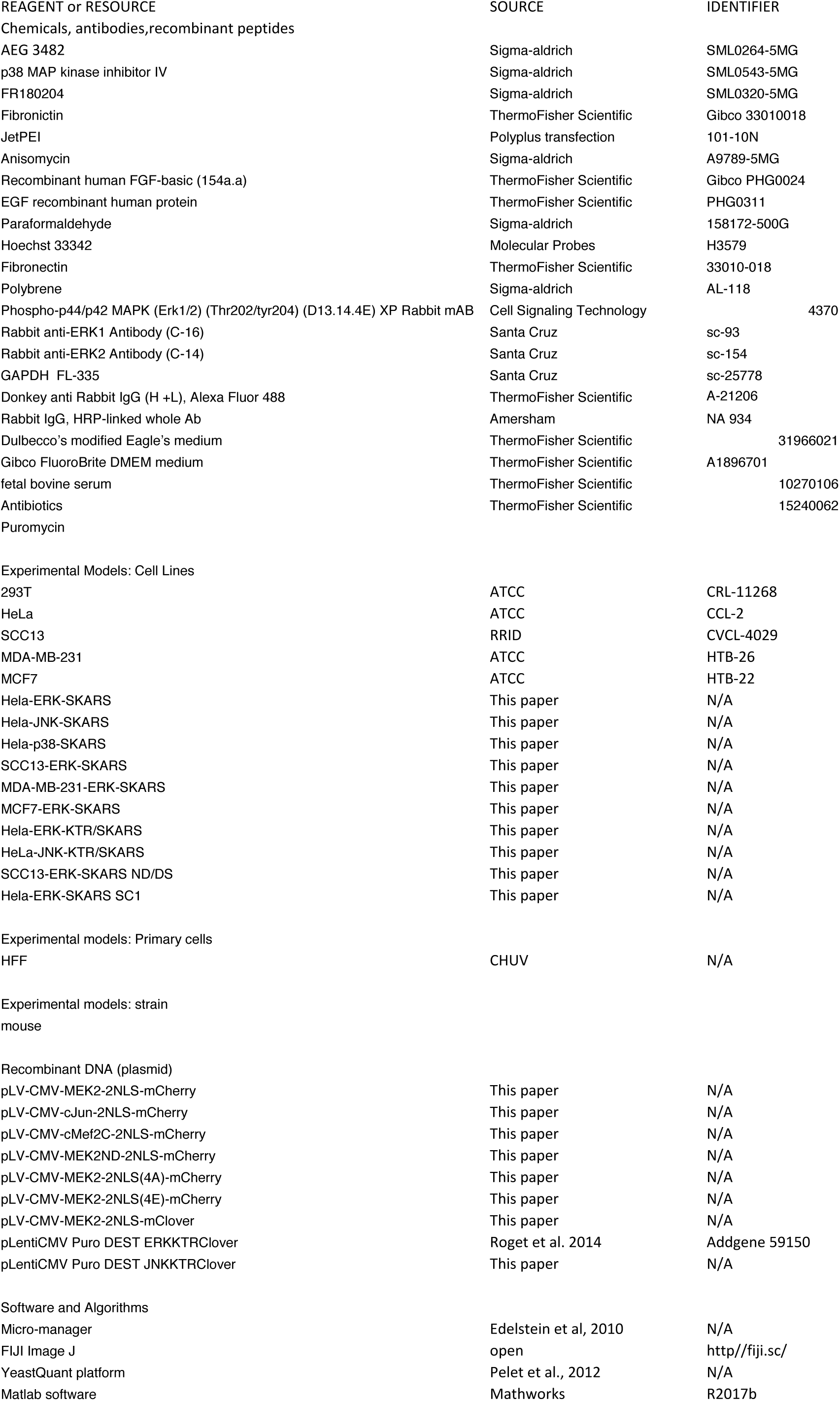

**Supplementary Movie 1. Pulses of ERK activity**

Time-lapse movie of HeLa cells expressing the ERK-SKARS displaying pulses in ERK activation upon 0.1 ng/ml EGF stimulation.

